# MobiChIP: a compatible library construction method of single-cell ChIP-seq based droplets

**DOI:** 10.1101/2023.12.31.573755

**Authors:** Xianhong Yu, Guantao Zheng, Liting Xu, Guodong Chen, Yiling Zhu, Tingting Li, Mingming Rao, Rong Cong, Wenshan Zheng, Hao Pei

**Author notes:** Correspondence (R.C.), (W.Z.), (H.P.) and (X.Y.). Contributed equally to this work (Xianhong Yu, Guantao Zheng, Liting Xu).

## Abstract

In order to illustrate the epigenetic heterogeneity, versatile tools of single-cell ChIP-seq (scChIP-seq) are necessary to meet the convenience and accuracy. Here, we develop MobiChIP, a compatible ChIP-seq library construction method based current sequencing platform with single cell level. As a novel capture strategy, MobiChIP is efficient to capture the fragments from tagmented nuclei of numerous species and execute the mixing of samples from different tissues or species. Especially, this strategy enables the flexible sequencing manipulation and sufficient nucleosome amplification without customized sequencing primers. MobiChIP reveals the landscape of chromatin regulation regions with active(H3K27ac) and repressive(H3K27me3) histone modification markers in peripheral blood mononuclear cells (PBMCs), and accurately unveiled the epigenetic repression of *hox* gene cluster in PBMCs than ATAC-seq. Meanwhile, we complete the bioinformatics pipeline to integrates the scChIP-seq data and scRNA-seq to illustrate the cellular epigenetic and genetic heterogeneity.

**One-Sentence Summary:** A high-throughput single-cell ChIP-seq based droplet reveals the integration of scRNA-seq data and scChIP-seq data.

---

Single-cell sequencing can demonstrate the complex heterogeneity of individual cells and reveal rare cell populations*(1-4)*. Single-cell technology has enabled transcriptomes sequencing(*5-6*), DNA methylation sequencing(*7-8*), DNase sequencing(*9-10*), assay for transposase accessible chromatin sequencing (ATAC-seq)(*11-14*). While ATAC and DNase can provide genome-wide snapshots of open chromatin, the study of diverse chromatin modifications and TF biding sites might provide further insights into epigenomic heterogeneity and cellular states. ChIP-seq, also known as a powerful tool for studying protein-DNA interactions in vivo and vitro, is usually used to reveal transcription factor binding sites or histone modification sites of genome(*15-18*). To unveil the relationship of gene expression and regulation, CoBATCH, CUT&Tag and ACT-seq based well plates and used protein A-Tn5(PATn5) have been reported to study protein-DNA interactions to reveal cell fate trajectories and population heterogeneities during developmental processes and pathological alterations in vivo and vitro, but are laborious(*19-24*). In order to obtain high-throughput of scChIP-seq, microfluidic Chip are developed to access scChIP-seq, while those methods belong to customized sequencing strategy(*25-26*).

Here, we develop a scChIP-seq method, MobiChIP, with a modified adaptor used to capture tagmented chromatin with protein A-Tn5. MobiChIP not need customized sequencing primers and is compatible with current sequencing platform in common use. Via MobiChIP, we investigated the epigenome of health PBMCs. The histone modification H3K27ac and H3K27me3 reveal the different landscape of active and repressive region of genome. We focused on the comparison of MobiChIP and ATAC-seq. Meanwhile, we integrated the epigenetic data with transcriptome to discrete heterogeneous populations. This study provides an experimental method and a bioinformatics pipeline to study epigenetic regulation and gene expression in complex tissues with a multi-omics dimension to obtain a deeper understanding of epigenetic heterogeneity in PBMCs.

## High-Throughput MobiChIP profiling of single cells

To perform MobiChIP on a large scale of cells to reveal the epigenetic heterogeneity, we combine the CoBATCH and microfluidic to generate the high-throughput single cell ChIP-seq based droplets (Fig. 1A). To enable the sequencing of scChIP-seq library agilely on current platform, we modified the capture adaptor of MobiChIP to sequence with pooled sample from others. We mixed equal number of human K562 cells and mouse L929 cells, and sequenced the resultant library. The data showed collision rate of ∼5.1%, suggesting a successful labeling of single cells (Fig. 1B). Total 350 mouse L929 cells and 469 human K562 cells were identified, clustered into two populations (Fig. 1C). The results showed that the mouse-derived cells and the human-derived cells were well separated, proving that captured signals of H3K27ac were useful for clustering analysis. Meanwhile, we further analyzed the number of valid fragments captured in different cells and found that the separation of cell clusters was independent to the number of captured valid fragments (Fig. 1D). In order to meet the sample mixing in some distinctive usage scenario, we also applied MobiChIP to map the histone modification with different samples in the same tube. Sample indexes of adaptor assembled into PATn5 were used to label different samples. Human K562, mouse 3T3 and hamster CHO and human were mixed to perform MobiChIP, and 97%, 96% and 95% labeling ratio was achieved respectively (Fig. 1E).

**Fig. 1.**
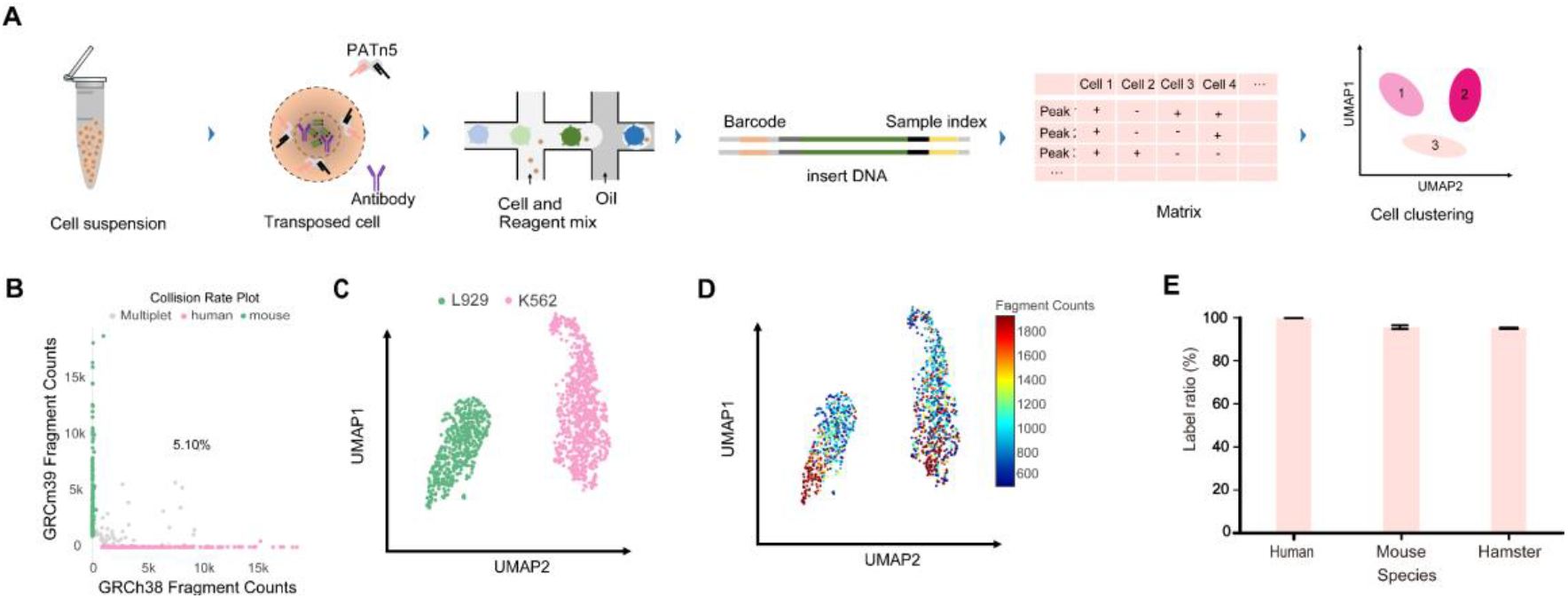
High-Throughput MobiChIP profling of single cells in different species. **(A)** Schematic of the MobiChIP workflow.This MobiChIP experiment contains the following procedure: sample preparation, ChIP and tagmentation, encapsulation, library preparation, sequencing and data analysis. **(B)** Scatterplots showing collision rate 5.10% for each unique barcode combination. Mouse L929 and human K562 cells were mixed at 1:1. **(C)** MAP projection of cells from (B) colored by cell clustering **(D)** MAP projection of cells from (B) colored by valid fragments. **(E)** The label ratio of human, mouse and hamster cells with barcoded PATn5 transposome.

## Single-cell profiles of several histone modifications in human samples

To validate the performance of MobiChIP, we performed experiments in several human samples. We identified single cells based on the number of fragments and fraction of fragments falling into peak regions, called from merged bulk data. Altogether, we performed MobiChIP with various histone modifications in K562 cell (H3K4me3) and health PBMCs (H3K4me1, H3K27ac, H3K27me3) respectively. Altogether, we obtained MobiChIP profiles of various histone modifications for 15,104 single cells, with the median unique fragments per cell: 1024 (H3K27ac), 1553 (H3K4me3), 1086 (H3K27me3) and 1457 (H3K4me1) (Fig. 2A). Total 25% to 58% of fragments fell within narrow peak regions, indicating a low level of background (Fig. 2B). The fragment length distribution was consistent with the capture of nucleosome fragments, as well as mono-, di- and tri -nucleosomes for all modifications (Fig. 2C).

**Fig. 2.**
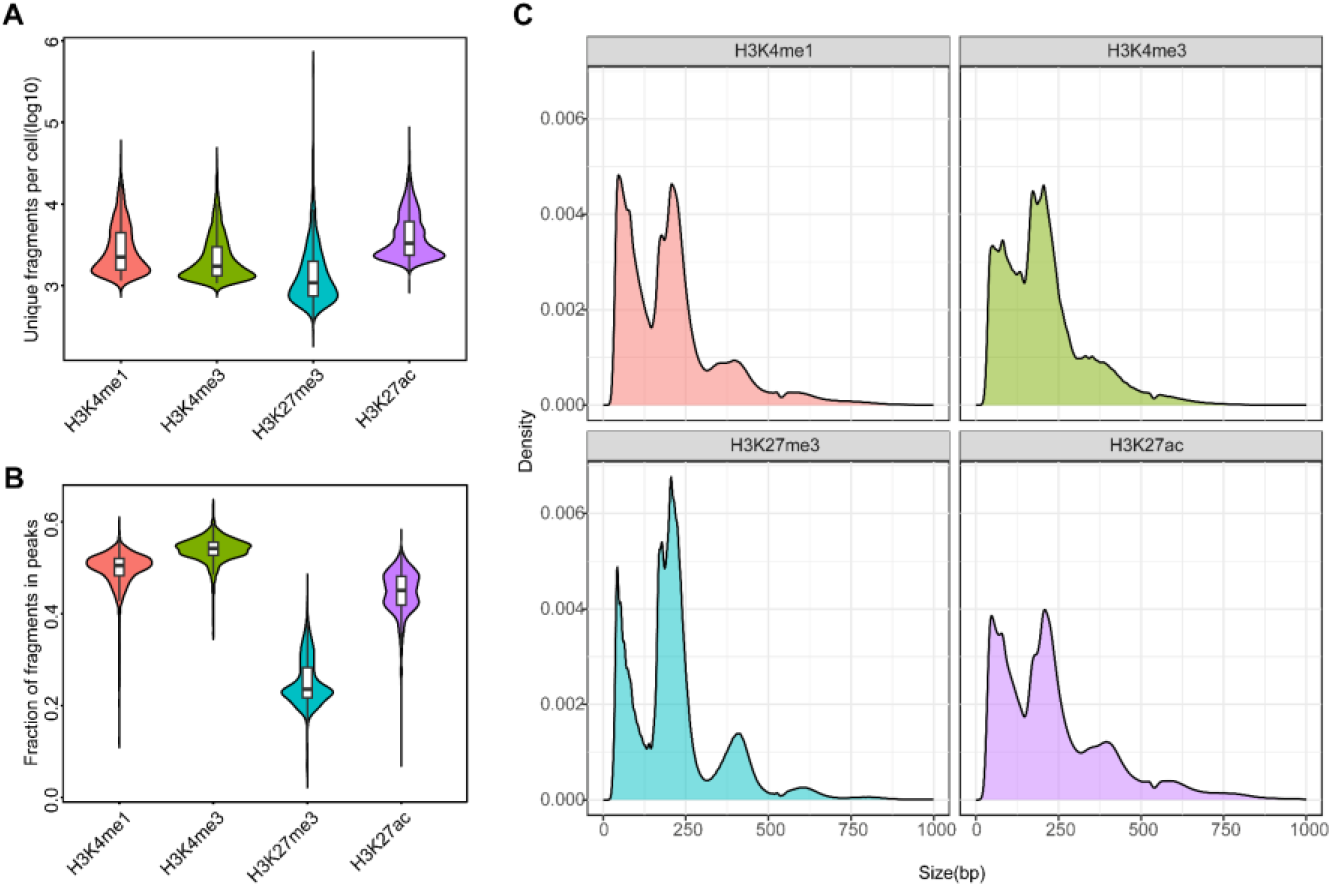
Single-cell profiles of several histone modifications in human samples. **(A)** Comparison of number of unique fragments per antibody per cell in MobiChIP. H3K4me1: n = 3,522 cells; H3K4me3: n = 1,918 cells; H3K27me3: n = 4,534 cells; H3K27ac: n = 5,130 cells. **(B)** Comparison of percentage of fragments falling into peak regions per antibody per cell in MobiChIP. Peaks were obtained by peak calling in merged bulk datasets. Cell number is same as in(A). **(C)** Distribution of fragment lengths in MobiChIP per antibody.

## Profiling of H3K27ac and H3K27me3 in single cells

To verify the fidelity of the MobiChIP, we mixed all single cells incubated with H3K27me3 antibody and compared the result with public data. By using the IGV snapshots, we found that the merged single-cell data showed a high consistency with the public data at the location of *Hox* gene cluster (Fig. 3A). We aggregated 500 single cells randomly sorted from PBMCs, the distribution of H3K27me3 signal was consistent with the aggregate one at *Hox* gene cluster region (Fig. 3A).

**Fig. 3.**
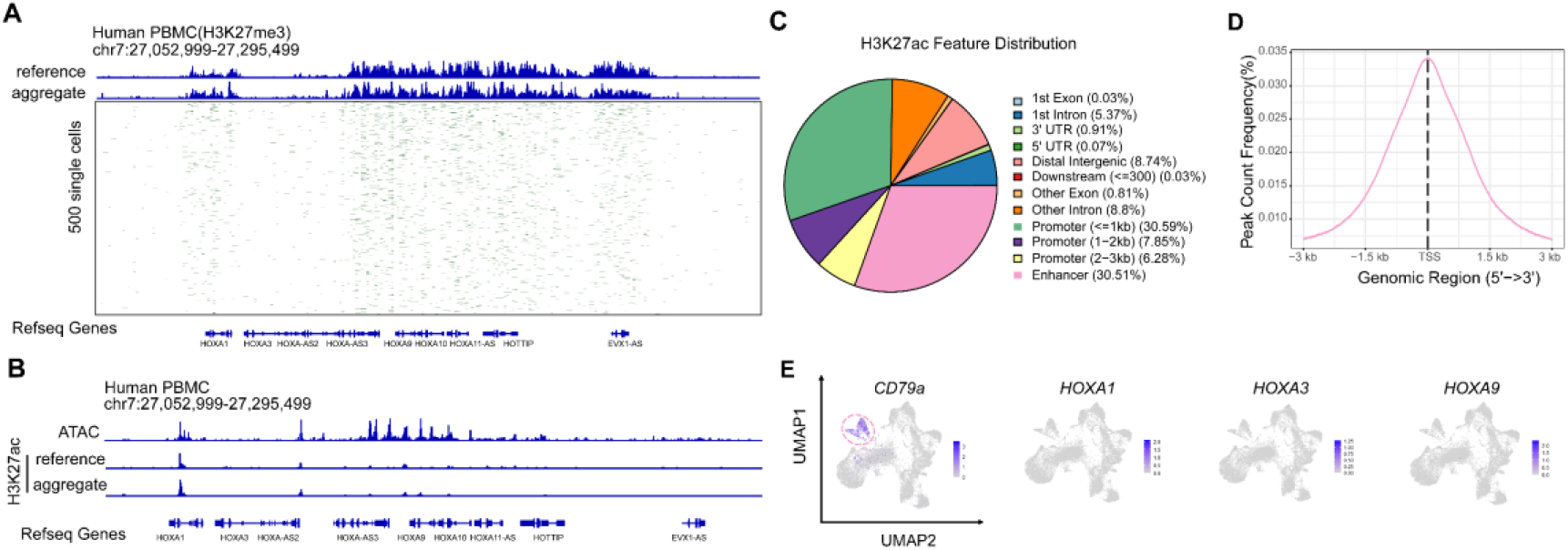
Profiling of H3K27ac and H3K27me3 in single cells. **(A)** Track plot representation of the MobiChP signal for H3K27me3 in PBMCs. The reference was downloaded from published data with H3K27me3. The x axis represents genomic region, each row in the y axis contains data from one cell. Signal is aggregated per 500 bp windows and binarized **(B)** Comparison of the MobiChP signal for H3K27ac with ATAC-seq signal. The reference(H3K27ac) and ATAC-seq was downloaded from published data. Peaks were obtained by peak calling in merged bulk datasets. **(C)** Distribution of H3K27ac feature at different elements of genome. **(D)** Peak count frequency of H3K27ac feature at TSS±3 kb. **(E)** Feature plot of *CD79a, H0XA1, HOXA3*, and *H0Xa9* genes in map projection of PBMCs with scRNA-seq. B cells were colored by red ring dashed line.

Meanwhile, the consistence suggested the quality of captured fragments from single cell was qualified to perform bioinformatic analysis. Since the histone modification H3K27me3 is a repressive epigenetic marker, the data suggests that gene expression in the *Hox* gene cluster is suppressed in PBMC cells. As a key gene for somite development, the expression of *Hox* family is significantly altered during blood cells mature. To demonstrate the epigenetic signal around the *Hox* gene cluster, we also performed MobiChIP experiments with H3K27ac antibody in PBMCs (Fig. 3B). We also accessed the H3K27ac ChIP-seq data as well as the ATAC-seq data from the public databases. Via the comparison we found that the ATAC-seq data indicated more chromatin open regions around the *Hox* gene cluster, while the MobiChIP and public ChIP-seq data indicated that the signal of H3K27ac around the *Hox* gene cluster was weak at a low level. In addition to that, site-specific analysis showed that the signal of H3K27ac was mainly present at promoter and enhancer regions (Fig. 3C), and the signals in TSS region were consistent with reported one (Fig. 3D). By combining with single-cell RNA-seq data, we found that clusters were specific and the native marker CD79A was expressed normally in B-cells, but *HOXA1, HOXA3* and *HOXA9* genes were not detected significantly in all cell types (Fig. 3E). Based on the above data, it is easy to conclude that two epigenetic modifications, H3K27me3 and H3K27ac, play an important role in suppressing the expression of related genes around the *Hox* gene cluster of PBMCs in blood, and also proves that MobiChIP is a more accurate tool to reveal the regulation of genes expression and cell fate decision than ATAC-seq.

## MobiChIP reveals epigenetic heterogeneity of health PBMCs

We performed histone modification of H3K4me1 in healthy human PBMCs and obtained total 10,031 cells and 1,467 unique DNA fragments per cell. By ArchR clustering and dimension reduction, we obtained five distinct cell populations, including B cells, CD4-positive T cells, CD8-positive T cells, NK cells and monocytes (Fig. 4A). In order to validate that the data quality of MobiChIP is reliable, we projected MobiChIP data with bulk ChIP-seq data (Fig. 4A). We accessed bulk ChIP-seq data for each cell type in PBMC cells from the database. Via mapping analysis, we found that the cell type annotated from the MobiChIP data corresponded well to the public data without bias. Furthermore, we found the annotation results of subpopulations were consistent with the projection of published single-cell RNA-seq data (Fig. 4B). Since single-cell ChIP-seq is used to study cellular heterogeneity at the level of gene regulation, we performed single-cell RNA-seq sequencing on this sample in order to visualize the combination of RNA expression information and DNA regulation information, and to make multi-omics comparisons of the cells at different dimensions. Single-cell RNA-seq sequencing yielded 14,453 cells with a median 1,889 genes per cell. By integrating and analyzing the data from single-cell ChIP-seq and single-cell RNA-seq through Signac, we found that the multi-omics data containing two different dimensions of information can be perfectly matched together (Fig. 4C). Through further subpopulations annotation, those data corroborated one another (Fig. 4D). Ultimately, our results showed that data analysis of MobiChIP can be unified with data from bulk ChIP-seq and single-cell RNA-seq, and integrated with single-cell RNA-seq. This strategy provides a powerful tool for us to be able to parse cellular epistasis and gene expression heterogeneity from different dimensions.

**Fig. 4.**
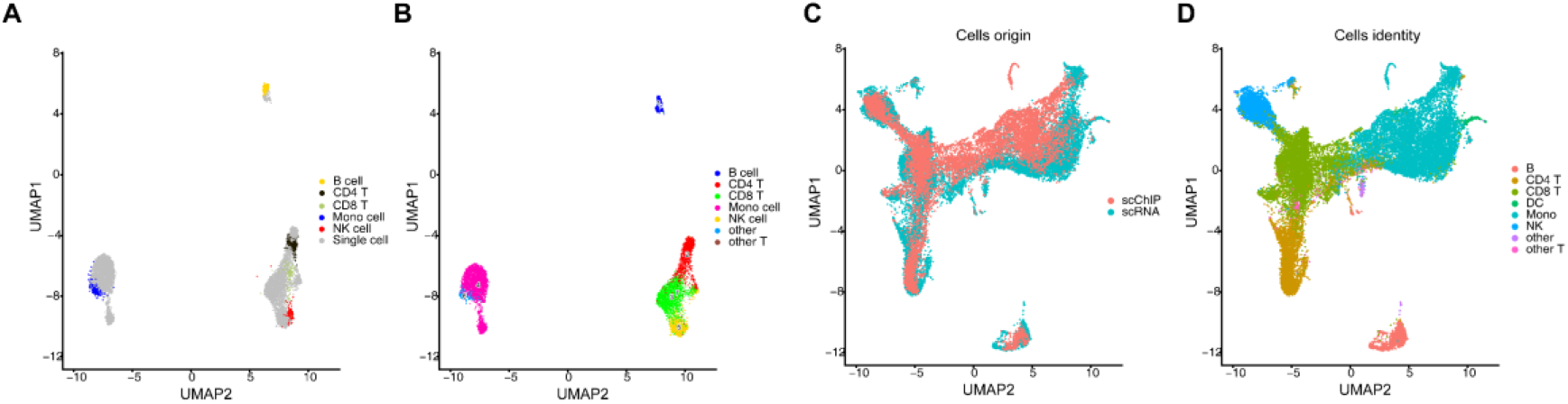
MobiChIP reveals epigenetic heterogeneity of health PBMCs. **(A)** Single cell ChlP-seq data of MobiChIP was projected with bulk ChIP data in terms of H3M4me1 in PBMCs. **(B)** Single cell ChlP-seq data of MobiChIP was projected with single cell RNA-seq data of PBMCs. **(C)** Co-embedding of scChlP-seq data and scRNA-seq data. **(D)** The integration and annotation of scRNA-seq data and scChlP-seq data.

## Discussion

We developed a new sequencing technology that is seamlessly compatible with existing sequencing platforms and facilitates users without customized sequencing primers. It also enables the mixing of samples from different species, which greatly reduces experimental costs and increases throughput of cells.

Cell heterogeneity of different tissues and organs plays an important role in disease development and progression. However, the relationship of gene expression and gene regulation relies on the specific ChIP-seq technology to accurately explain. Via performing MobiChIP and RNA-seq on PBMCs, we identified a strong signal for H3K27me3 (a repressive marker) around the *Hox* gene cluster in PBMCs, while the signal of H3K27ac (an active marker) was weak at a low level, suggesting that the alteration of epigenetic modification plays an important role in the maturation process of PBMCs. We also fully investigated the data of ATAC-seq, and the chromatin accessibility still remains around the *Hox* gene cluster, suggesting the ATAC-seq data not corresponded with the repression of *Hox* gene. Therefore, MobiChIP has a higher precision and can distinguish the differences in cellular epigenetic heterogeneity. Meanwhile, combining the scRNA-seq data demonstrated that the genes in the *Hox* gene cluster, whose expression is gradually closed with the differentiation of blood cells.

Since cell fate decisions are induced by the regulation of multiple dimensions, recently multi-omics technologies have also been developed. In order to enable the MobiChIP technology more applicable, we also build a set of process for multi-omics analysis. By performing MobiChIP and scRNA-seq sequencing with the same sample of PBMCs, we found that the data from MobiChIP can match well with the data from single-cell RNA-seq. The data from two dimensions can corroborate each other in terms of clustering and annotation. It also proves that the multi-omics analysis process we developed is fully capable of data union in multiple dimensions.

The limitation of MobiChIP platform is insufficient depth of bioinformatics analysis for the time being. Using MobiChIP to unveil the development trajectory, cell specific regulatory elements and some new regulatory targets need be further studied. However, as a versatile tool, we foresee that MobiChIP would help to get more biological information from DNA regulation level, to reveal epigenetic and genetic heterogeneity, coupled with 3’ transcriptome, 5’ transcriptome, VDJ and MobiCITE of MobiDrop single cell sequencing technology. We also hope that researchers using this technology together to promote this platform to further improve the experiment and bioinformatics analysis process.

## Supporting information

Supplementary materials will be used for the link to the file on the preprint site.

## Acknowledgments

We thank all members of MobiDrop for critical comments on this manuscript. We thank the Mingma Technologies (Shanghai, China) for help of next sequencing.

## Funding

X.Y. and H.P. were supported by the Microfluidic Bioassay Technology Venture Team Program of China (2022R02005), the Automatic Digital PCR All-in-One Machine Program of China (2021C03199).

## Author contributions

X.Y., H.P., W.Z. and R.C. conceived and designed the study. X.Y. designed and performed all experiments. G.Z. and L.X. performed the computational analyses. G.C. provided technical support supervised by X.Y. X.Y. wrote the paper with input from all other authors. All authors participated in data discussion and interpretation.

## Competing interests

The authors declare no competing interests.

## Notes

### Competing Interest Statement

The authors have declared no competing interest.

