## Supplementary materials will be used for the link to the file on the preprint site. for "MobiChIP: a compatible library construction method of single-cell ChIP-seq based droplets"

**Materials and Methods**

**Cell culture**

Human K562 cells were cultured in IMEM culture medium containing 1% fetal bovine serum (Sigma), 1% Penicillin/Streptomycin (Hyclone) at 37°C with 5% $CO_2$. Mouse L-929 cells were cultured in MEM culture medium containing 1% fetal bovine serum (Sigma), 1% Penicillin/Streptomycin (Hyclone) at 37°C with 5% $CO_2$. Hamster CHO cells were cultured in RPMI culture medium containing 1% fetal bovine serum (Sigma), 1% Penicillin/Streptomycin (Hyclone) at 37°C with 5% $CO_2$.

**Human PBMCs and mouse primary cells**

Frozen healthy human PBMCs and mouse primary cells from spleen(C57BL/6) were purchased from ORIBIOTECH. Frozen primary cells were thawed according to the manufacturer's instructions.

**Antibodies**

Antibodies used were H3K4me3 (1:100, Diagenode, C15410030), H3K27ac (1:100, Abcam, Ab177178), H3K27me3 (1:100, Active Motifs, 39055), H3K4me1 (1:100, Abcam, ab8895) and guinea pig anti-rabbit (1:50, ABIN1011961).

**PATn5 transposome assembly**

PATn5 transposome was purchased from Active Motifs (Cat. NO. 53161). Assembly of PATn5 transposome referred to the manufacturer's instructions. Oligonucleotides (Table S1) were dissolved in buffer (10 mM Tris, pH 8.0) to make 100 μM stock solution. To anneal adaptors, MEA Oligo, MED Oligo, barcoded Oligo 1, barcoded Oligo 2, or barcoded Oligo 3, were mixed with equal volume of MErev oligo respectively, placed at a thermal cycler for 5 min at 92 °C followed by programmed temperature decrease at 0.1 °C /s to 20 °C, and kept in 8 °C. To get 4 mM assembled PATn5 transposome complex, 2 mM annealed adaptor MEA was mixed with 2 mM annealed adaptor MED, barcoded adaptor 1, barcoded adaptor 2, or barcoded adaptor 3 respectively, 4 mM transposase PATn5 and storage buffer (100 mM HEPES pH7.2, 200 mM NaCl, 0.2 mM EDTA, 2 mM DTT, 0.2% Triton X-100, 50% glycerol) and incubating at room temperature for 50 min. The assembled PATn5 transposome was stored -20 °C.

**Activity measurement of PATn5 transposome**

The activity of assembled PATn5 transposome was assayed by tagmentation of human genomic DNA as previously described (Chenet al., 2016; Picelli et al., 2014; Wang et al., 2019). 1 μl 4 mM barcoded PATn5 transposome was mixed with 100 ng human genomic DNA and 2 μl TAPS-MgCl2-DMF (50 mM TAPS-KOH pH 8.3, 50 mM MgCl2, 50% DMF) in 10 μl system. The tagmentation was incubated at 55 °C for 10 min and stopped by adding 2 μl Stopping Buffer (250 mM EDTA, 0.2% SDS) to incubation for 10 min. The fragmented DNA was resolved on 1.5% agarose gel for examination of size distribution.

**MobiChIP procedure**

MobiChIP was performed as in CoBATCH with minor modification described below. The MobiChIP was performed in 200 μl tubes, all washes and incubation volumes were 200 μl. All centrifugations were done using a swinging bucket centrifuge with an adapter. Total 200,000 to 500,000 cells were thrown directly into a 200 μl tube and centrifuged at 4 °C for 3 min at 300 g. The cell pellet was resuspended with 100 μl of antibody buffer (20 mM HEPES pH 7.6, 150 mM NaCl, 2 mM EDTA, 0.5 mM spermidine, 0.05% digitonin, 0.01 % Triton X-100, 1x protease inhibitors) containing a primary antibody and incubated at 4 °C for 2-3 h. Following the incubation, the tube was centrifuged at 4 °C for 3 min at 300 g and nuclei pellet was resuspended with 100 μl of antibody buffer with 1:50 diluted secondary antibody. After the incubation for 20 min, the nuclei were centrifuged at 4 °C for 3 min at 300g, washed once with 200 μl of Dig-wash buffer (20 mM HEPES pH 7.6, 150 mM NaCl, 0.5 mM spermidine, 0.05% digitonin, 0.01% Triton X-100, 1x protease inhibitors), resuspended in 200 μl of Dig-wash buffer with 1:100 diluted protein A-Tn5 fusion and incubated for 1 h rotating at 4 °C. Then, nuclei were centrifuged for 3 min at 300g, washed two times with 200 μl of Dig-wash buffer, resuspended in 15 μl tagmentation buffer (10 mM TAPS-NaOH pH 8.3, 10 mM MgCl2) and incubated for 45 min at 37 °C. Following tagmentation, 135 μl PBS containing 0.1% BSA was added by gently pipetting up and down several times. The nuclei were centrifuged for 3 min at 300g and resuspended with 50 μl PBS containing 0.1% BSA.

**Barcoded MobiChIP procedure**

Barcoded MobiChIP procedure referred to MobiChIP procedure described above. K562 cells, L-929 cells and SHZ-88 cells were pipeted directly into a 200 μl tube respectively, and following independent manipulation. Barcoded PATn5 transposomes were used to distinguish those cells from different species. At the end of tagmentation, the nuclei of K562 cells, L-929 cells and CHO cells were retrieved with centrifugation at 300g for 3 min and resuspended with PBS containing 0.1% BSA. Those nuclei were counted and combined with equal amount.

**MobiChIP library preparation and sequencing**

The single nucleus suspension was loaded into microfluidic chip of ChIP C Single Cell Kit (MobiDrop (Zhejiang) Co., Ltd., cat. no. S190200101) to obtain droplets with MobiNova-100 (MobiDrop (Zhejiang) Co., Ltd., cat. no. A1A40001). Each nucleus was wrapped into a droplet which contained amplification reagent and a gel bead linked with up to millions oligos (cell unique barcode). After encapsulation, droplets suffer light cut by MobiNovaSP-100(MobiDrop (Zhejiang)

Co., Ltd., cat. no. A2A40001) following oligos diffusion into amplification mix. The tagmented chromatin DNA was captured by gel beads containing capture sequence in droplets. Following preamplification, linearized DNA with barcodes were amplified, and a library was constructed using the High Throughput Single Cell ChIP-seq Kit (MobiDrop (Zhejiang) Co., Ltd., cat. no. S190300101) and the ChIP-seq Dual Index Kit (MobiDrop (Zhejiang) Co., Ltd., cat. no. S190400101). The MobiChIP libraries were sequenced on an Illumina NovaSeq 6000 sequencing system (paired-end 150bp) by Mingma Technologies (Shanghai, China).

**scRNA-seq library preparation and sequencing**

The single cell suspension was loaded into microfluidic chip of Chip A Single Cell Kit v2.0 (MobiDrop (Zhejiang) Co., Ltd., cat. no. S050100201) to generate droplets with MobiNova-100(MobiDrop (Zhejiang) Co., Ltd., cat. no. A1A40001). Each cell was wrapped into a droplet which contained reaction reagent and a gel bead linked with up to millions oligos (cell unique barcode). After encapsulation, droplets suffer light cut by MobiNovaSP-100(MobiDrop (Zhejiang) Co., Ltd., cat. no. A2A40001) following oligos diffusion into reaction mix. The mRNAs were captured by gel beads with cDNA amplification in droplets. Following reverse transcription, cDNAs with barcodes were amplified, and a library was constructed using the High Throughput Single Cell 3’RNA-Seq Kit v2.0 (MobiDrop (Zhejiang) Co., Ltd., cat. no. S050200201) and the 3' Single Index Kit (MobiDrop (Zhejiang) Co., Ltd., cat. no. S050300201). The scRNA-seq libraries were sequenced on an Illumina NovaSeq 6000 sequencing system (paired-end 150bp) by Mingma Technologies (Shanghai, China).

**scRNA-seq data process**

Raw datas (fastq format) of single cell transcriptomic were pre-analyzed by MobiVision v1.1(MobiDrop), and reads were aligned to Homo sapiens reference GRCh38 and Mus musculus reference GRCm39. Filtered cell-gene matrix was obtained with MobiVision v1.1. For further analysis, low-quality cells were filtered out according to the methods of disclosure(*1*).

**Barcoded MobiChIP data process**

Paired-end sequencing reads from MobiChIP libraries with Illumina sequencing platform (fastq format) were processed using the pipeline called MobiVision-v3.0 and integrated multi-omics analysis (https://www.mobidrop.com/bioinformatics-analysis-software/mobivision-news/software-download). QC report and other relative result files were obtained. The pipeline was generated by five steps: barcodes correction, reads trimming, alignment, peaks calling and cell calling. Sequences with mismatch≤2 bp were allowed in barcodes correction while the useless sequencing adaptors were removed. The passed reads were aligned to the human (GRCh38), mouse (GRCm39) or hamster (criGriChoV2) reference genome with Bowtie2(*2*) in alignment step. Removing duplications was followed and peaks were called with MACS2(*3*). Cell-associated-barcodes were filtered with the existing method published by Zheng et al.(*4*). All barcodes and fragments relative files would be output.

In this study, the filtered fragment data with cell barcodes were first encapsulated into a BED file. This file was subsequently uploaded into ArchR(*5*), facilitating fragment count computations within 5-kb genomic windows. This step was integral to all dimensionality reduction processes in our experiments. For dimensionality reduction, we employed Latent Semantic Indexing (LSI) with a Term Frequency-Inverse Document Frequency (TFIDF) normalization approach(*6*). Additionally, we utilized Uniform Manifold Approximation and Projection (UMAP) for low-dimensional embedding and executed clustering based on a nearest-neighbor graph in the LSI space.

To identify cell types accurately, our process began with using ArchR to compute gene activity scores, which reflect the activity levels of genes. We then identified genes with high scores in each cluster. These high-scoring genes were matched with published cell-type markers to accurately categorize each cluster.

**Integrating MobiChIP data with scRNA-seq**

In integrating single-cell transcriptome data with single-cell ChIP-seq data, we utilized ArchR's 'addGeneIntegrationMatrix' feature. The integration mechanism aligns cells from scChIP-seq with those from scRNA-seq by comparing the gene score matrix of the former with the gene expression matrix of the latter. For each scChIP-seq cell, the integration process identifies the most similar cell in the scRNA-seq dataset, assigning the cell type information from this scRNA-seq cell to the scATAC-seq cell. Consequently, every cell in the scChIP-seq dataset acquires a gene expression signature, paving the way for various downstream analyses.
